# Loss of *Tbx4* Affects Postnatal Lung Development and Predisposes to Pulmonary Hypertension

**DOI:** 10.1101/2024.09.18.613783

**Authors:** Gabriel Maldonado-Velez, Elizabeth A. Mickler, Todd G. Cook, Micheala A. Aldred

## Abstract

Pulmonary arterial hypertension (PAH) is a progressive vascular disease characterized by remodeling of the precapillary pulmonary arteries. Genomic variation within the T-box 4 (TBX4) transcription factor is the second most common genetic cause of PAH, and can also cause severe lung developmental disorders with neonatal PH. Currently, the effect of TBX4 loss-of-function on later stages of lung development and predisposition to lung disease, including PH, is not well understood. Therefore, we have generated *Tbx4* conditional knockout (*Tbx4-CKO*) mice in which *Cre recombinase* deletes exon 5 of *Tbx4* within the embryonic lung mesenchyme to create a null allele. We harvested lungs from these mice at various timepoints to examine alveologenesis, vascularization, vascular remodeling, lung cellular composition, and disruption of transcriptional activity compared with control lungs. Right ventricular systolic pressure (RVSP) was measured in six-month-old mice to evaluate for PH. *Tbx4-CKO* lungs show enlargement of airspaces, as confirmed by an increase in mean linear intercept at P14 (24.9%), P36 (31.5%), and P180 (49.6%). These lungs also show a 39.3% decrease in von Willebrand Factor-positive vessels and a 14.2% increase in vessel wall thickness. Consistent with these results, *Tbx4-CKO* mice show a statistically significant increase of 15.7% in RVSP and 16.3% in the Fulton index. Bulk-RNA sequencing analysis revealed enrichment of pathways and genes relevant to lung alveologenesis, angiogenesis, and PH. Our results show that disruption of *Tbx4* expression during early lung development is sufficient to disrupt postnatal lung development and circulation.

## Introduction

The term pulmonary hypertension describes a group of vascular disorders of the cardiopulmonary system characterized by a mean pulmonary artery pressure (mPAP) >20 mmHg at rest. The World Symposium on Pulmonary Hypertension (WSPH) divides PH into five different categories based on the etiology of the disease. Group 1 PH, also known as pulmonary arterial hypertension (PAH), is a rare disease affecting 15 to 50 individuals per million and involves endothelial dysfunction, progressive vascular remodeling, and loss of small pulmonary arteries [1, 2]. Genetic variation within several genes is known to contribute to the etiology of PAH, but approximately 70-80% of the heritable form of PAH cases can be explained by mutations within the bone morphogenetic protein receptor 2 (*BMPR2*) [3]. Group 3 PH is the second most common cause of PH and is characterized by complications of lung diseases, including chronic obstructive pulmonary disease (COPD), diffuse parenchymal lung disease, hypoxia, and developmental lung disorders [4]. Histological examination of lungs with COPD and idiopathic pulmonary fibrosis (IPF) indicate that loss of small vessels, reduction of capillary bed due to destruction of alveolar septa, compression of the vasculature due to accumulation of collagen fibers, and vascular remodeling are part of the histopathologic changes in Group 3 PH [4]. Despite current advances in knowledge and therapies, more studies are required to understand the molecular mechanism underlying these pulmonary vascular disorders and help improve diagnosis and current treatment outcomes.

The TBX4 protein is a transcription factor robustly expressed, among other tissues, within the embryonic mouse hindlimb, genital tubercle, trachea, and lung mesenchyme [5, 6]. *Tbx4*-expressing mesenchymal cells give rise to multiple cell lineages within the developing lung, suggesting that it plays an essential role in various aspects of lung development [7]. Constitutive inactivation of *Tbx4* expression reduces branching morphogenesis in E13.5 mouse lungs [5]. Consistent with these results, a recent study reported a reduction in the number of branching tips in mouse embryonic lung explants after knocking down *Tbx4* [8]. In humans, variants within the *TBX4* genomic locus are the second most common genetic cause of PAH and are mainly associated with pediatric onset of PAH when compared to variants in other risk genes [9, 10]. Infants with *TBX4* deleterious variants may also display lethal lung developmental disorders, including acinar dysplasia (AcDys) and congenital alveolar dysplasia (CAD), and develop PH [11]. Thus, *TBX4* variants in humans are associated with significant clinical heterogeneity at the histological and physiological levels. The overlapping of pathologic features present challenges to make an accurate diagnosis. Therefore, more studies are needed to characterize better the effect of TBX4 loss of function (LoF) in lung development and how developmental abnormalities contribute to disease, including PH.

In this study, using mouse genetics, we disrupted *Tbx4* expression in the lung mesenchymal lineage by conditionally deleting exon 5, which encodes a portion of its DNA-binding domain. Our results show that conditional knockout of *Tbx4* results in abnormal postnatal lung development and predisposition to pulmonary hypertension, similar to observations made in human cases with *TBX4* mutations. Some of the results of these studies have been previously reported in the form of abstracts [12, 13].

## Methods

Detailed methods are provided in the online data supplement. Briefly, *Tbx4^LME^-Cre* (IMSR_JAX:033331) and *Tbx4^cond^* (MMRRC_043812-JAX) mouse models were purchased from The Jackson Laboratory [14, 15] and crossed to generate homozygous conditional knockouts (*Tbx4-CKO*). All animal maintenance and procedures were performed in accordance with the Animal Care and Use Committee at Indiana University School of Medicine. After euthanasia, murine lungs were perfused with cold PBS through the right ventricle to flush out the blood. Mice at various postnatal stages were euthanized and lungs were inflated by instillation with 10% neutral buffered formalin (NBF) using a tracheal cannula at a constant pressure of 20 or 25 cmH2O [16]. Fixed lungs were used to measure multiple parameters, including mean linear intercept (MLI), number of vessels per high power field (HPF), and vessel wall thickness. The MLI was measured from hematoxylin and eosin (H&E) stained tissues sections at various time points [17]. To evaluate pulmonary vascularization, lung sections were immunostained with the endothelial cell marker, von Willebrand factor (vWF). Subsequently, we counted the vWF-positive vessels with a diameter less than 100 μm per HPF [18]. Masson’s Trichrome and Verhoeff’s Van Gieson stains were used to visualize elastin in the blood vessels and measure the vessel wall thickness. Echocardiography was performed using a Vevo 2100 ultrasound machine (VisualSonic) with a 40 MHz probe. The Vevo Lab software was used to measure diastolic and systolic dimensions, RVOT, TAPSE, cardiac output, stroke volume, pulmonary acceleration time (PAT), and pulmonary ejection time (PET). To measure right ventricular systolic pressure (RVSP), a midline neck incision was made and the right external jugular vein was cannulated with a 1.4F Millar catheter, which was advanced into the right ventricle. At the end of the procedure, the anesthetized mouse was sacrificed via exsanguination and excision of the heart. A previously described whole-mount or tissue sections RNA *in situ* hybridization (ISH) protocol was used to evaluate gene expression in mouse embryos [19, 20]. For adult lungs, the RNAscope Multiplex Fluorescent Reagents Kit v2 (Advanced Cell Diagnostics, Intro Pack, Cat. No. 323136) was used following the manufacturer’s instructions. Total RNA isolation from *Tbx4-CKO* (n = 3) and *Tbx4^fl/fl^* (n = 3) lungs was performed using the RNeasy Mini Kit (Qiagen). The changes in gene expression were investigated by quantitative reverse transcription polymerase chain reaction (RT-qPCR) analysis or RNAseq. For RNAseq, the RNA integrity number (RIN) for each sample was obtained using the 4200 TapeStation system (Agilent). Library preparation was performed using the KAPA RNA HyperPrep Kit (Roche) and total RNA. Paired-end sequencing was performed on an Illumina NovaSeq 6000 instrument. Differential expression analysis was performed, and genes with a false discovery rate (FDR) < 0.05 and fold-change > 2 were considered significant. Data will be available in GEO.

### Statistical Analysis

To assess the normality of the data, we used the Shapiro-Wilk test. Subsequently, a parametric unpaired two-tailed *t* test or nonparametric Mann-Whitney test was used to determine statistical significance, which was defined as *P* < 0.05. Descriptive statistics are shown throughout the text and indicate the mean ± SEM.

## Results

### Disruption of *Tbx4* Expression Within the Lung Mesenchyme of Mouse Embryos

Constitutive deletion of *Tbx4* causes embryonic lethality due to allantois developmental defects and failure of chorioallantoic fusion [5, 8, 21]. To circumvent this issue, we have generated *Tbx4* conditional knockout (*Tbx4-CKO*) mice by mating transgenic *Tbx4^LME^-Cre* and *Tbx4^cond^* mice [15, 22]. In *Tbx4-CKO* mice, the *Tbx4* lung enhancer controls Cre recombinase expression within the mesenchyme, where it deletes exon 5 of *Tbx4* (Figure 1A). This exon encodes a portion of the DNA-binding T-box domain of TBX4, and therefore, its removal is predicted to disrupt the function of the protein. In addition, Cre recombinase-mediated excision of exon 5 results in the formation of a premature stop codon at the junction of exon 4 and 6 of *Tbx4* (Figure E1). This may result in the degradation of the mutant transcripts through nonsense-mediated mRNA decay (NMD) or its translation into a truncated protein. To characterize our mouse model, we first confirmed the tissue-specific expression of Cre recombinase by RNA ISH using an antisense riboprobe. As expected, its expression pattern is restricted to the lung mesenchyme in the embryonic mouse lung (Figures 1B, 1C, and 1D). Subsequently, to ensure we are effectively disrupting *Tbx4* expression, we performed RNA ISH using an antisense riboprobe that spans exons 4, 5, and 6. The results show a marked reduction of *Tbx4* transcripts in the embryonic day 13.5 (E13.5) *Tbx4-CKO* lungs (Figure 1E). Consistent with this result, RT-qPCR analysis shows effective deletion of exon 5, as confirmed by over a 90% reduction in *Tbx4* wild-type (WT) transcripts within E13.5 *Tbx4-CKO* lungs (1.13 ± 0.25 versus 0.03 ± 0.01; *P* = 0.0018) (Figure 1F). Since we are only removing exon 5, we next sought to evaluate whether *Tbx4* transcripts lacking exon 5 are stable or undergo NMD. To this end, we designed primers to amplify different exons of the *Tbx4* transcript and measured expression levels. The RT-qPCR results show that the mutant transcripts undergo at least partial NMD (*Tbx4_*e4-e6: 1.04 ± 0.12 versus 0.25 ± 0.04; *P* = 0.0003 and *Tbx4_*e7-e8: 1.18 ± 0.26 versus 0.48 ± 0.06; *P* = 0.0387) (Figures 1G and 1H). Taken together, these results show that we are effectively disrupting *Tbx4* expression within the lung mesenchyme of mouse embryos.

**Figure 1.**
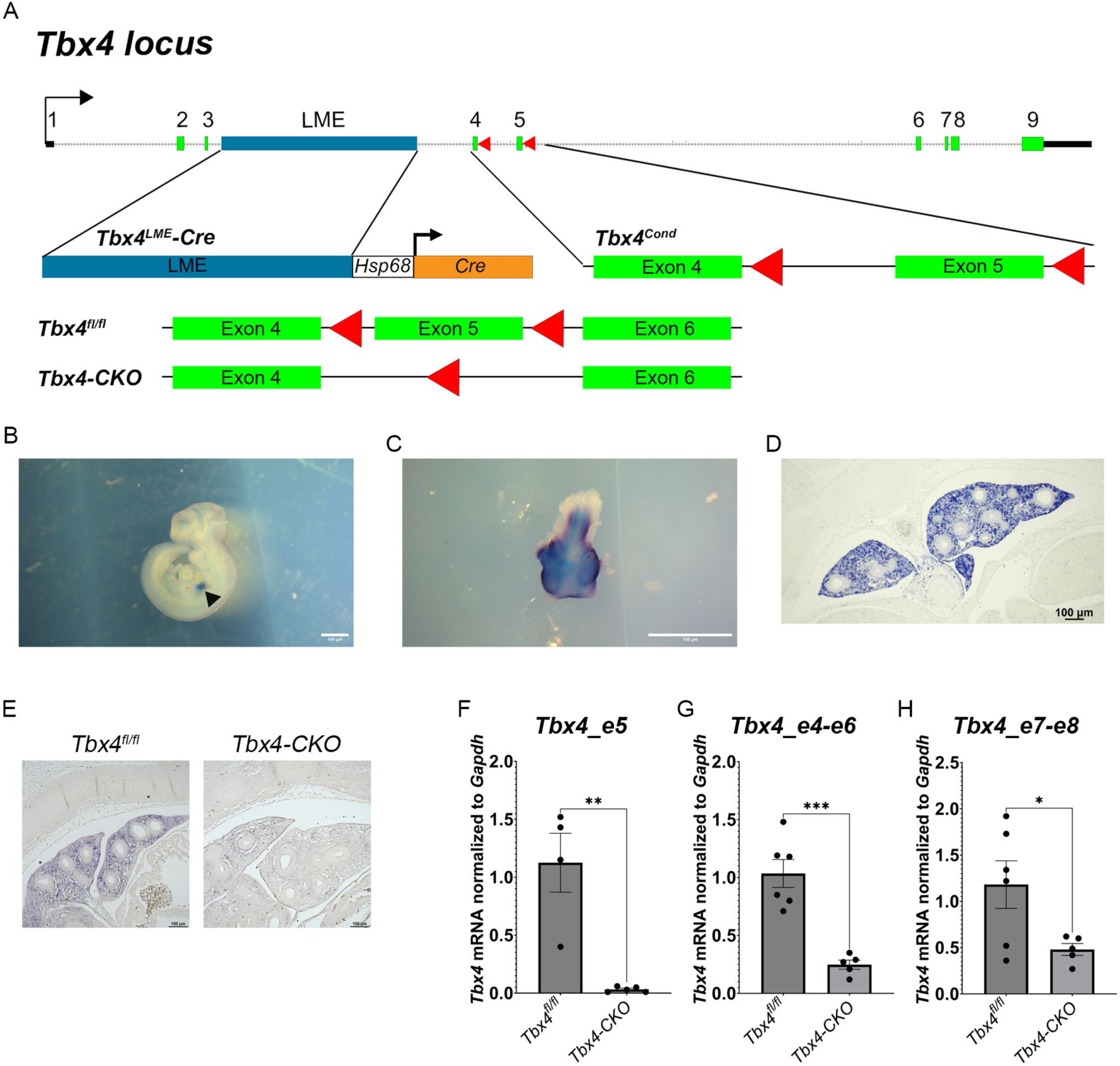
Effective disruption of *Tbx4* expression within the lung mesenchyme of mouse embryos. (A) Schematic showing the mouse *Tbx4* locus and the genetics of the mice used in this study. The LME (blue box) enhancer sequence within the *Tbx4* locus was used in the *Tbx4^LME^-Cre* construct to control expression of Cre Recombinase within the developing mouse lung. The *Tbx4^cond^*allele contains loxP sites (red arrowheads) flanking exon 5. The green boxes represent the exons, the black boxes represent the 5 and 3 prime untranslated regions, and the black arrow is the transcriptional start site. The control mice in the study are of the genotype *Tbx4^fl/fl^* and the *Tbx4-CKO* mice are of the genotype *Tbx4^LME^-Cre*; *Tbx4^fl/fl^*. (B) Whole-Mount ISH (WMISH) using Cre antisense riboprobes to validate expression of the Cre Recombinase. Black arrow points at robust staining in the lung of an E10.5 embryo. (C) WMISH using Cre antisense riboprobe on E11.5 lungs. (D) RNA ISH for Cre using lung tissue sections from E13.5 mouse embryos. (E) RNA ISH for *Tbx4* using lung tissue sections from E13.5 mouse embryos. (F-H) RT-qPCR results using sets of primers to amplify different exons of the *Tbx4* transcript. The scale bars are 100 μm. The statistical test used was a parametric unpaired two-tailed *t test*. \**P* < 0.05; \*\**P* < 0.01; \*\*\**P* < 0.001.

### Disruption of *Tbx4* Expression in Early Lung Development Affects Alveolarization

*Tbx4* expression is detected within the mesenchyme of the primitive lung in mice, starting approximately at E9.5 [5]. Its expression peaks at E15 and subsequently declines sharply by E18 [8]. The studies reporting that evidence have demonstrated that reducing *Tbx4* expression reduces branching morphogenesis and disrupts the expression of key transcriptional regulators within the embryonic mouse lung. The expression pattern of TBX4 suggests that it plays an essential role in developmental processes at that timepoint, including the extension of the epithelial tubes deeper into the mesenchyme and the formation of the future airspaces. Lung development is divided into multiple stages, and disruption of the early developmental processes can trigger a chain reaction effect that affects the subsequent stages. In mice, the alveolarization of the lungs is divided into two phases: classical, also known as bulk, and continued alveolarization [23]. To examine the effect of early disruption of *Tbx4* expression on this process, we harvested lungs during the bulk alveolarization at P14 and at the end of the process around P36. *Tbx4-CKO* mice are viable, good breeders, and seemed healthy based on body weight measurements at various timepoints (Figure E2). A simple histological examination revealed enlargement of airspaces, also known as alveolar simplification, at various timepoints in *Tbx4-CKO* lungs (Figure 2A). Alveolar simplification was confirmed by measuring the MLI, which shows a statistically significant increase of 24.9% (29.92 ± 1.11 versus 37.37 ± 1.29; *P* = 0.0014) and 31.5% (32.43 ± 1.65 versus 42.65 ± 1.80; *P* = 0.0018) at P14 and P36, respectively (Figure 2B). To examine the progression of this phenotype, we also harvested lungs from six-month-old mice. The MLI from adult lungs shows a more significant increase of 49.6% (34.84 ± 0.60 versus 52.16 ± 2.46; *P* < 0.0001) in the *Tbx4-CKO* group. In addition to alveolar simplification, closer examination of the lung tissue sections revealed regions of thickened and fibrous parenchyma, which are pathologic features observed in human cases with *TBX4* mutations (Figure E3). In addition, one of the mutant lungs contains organized thrombi occluding two small pulmonary vessels, which is also observed in PH (Figure E3).

**Figure 2.**
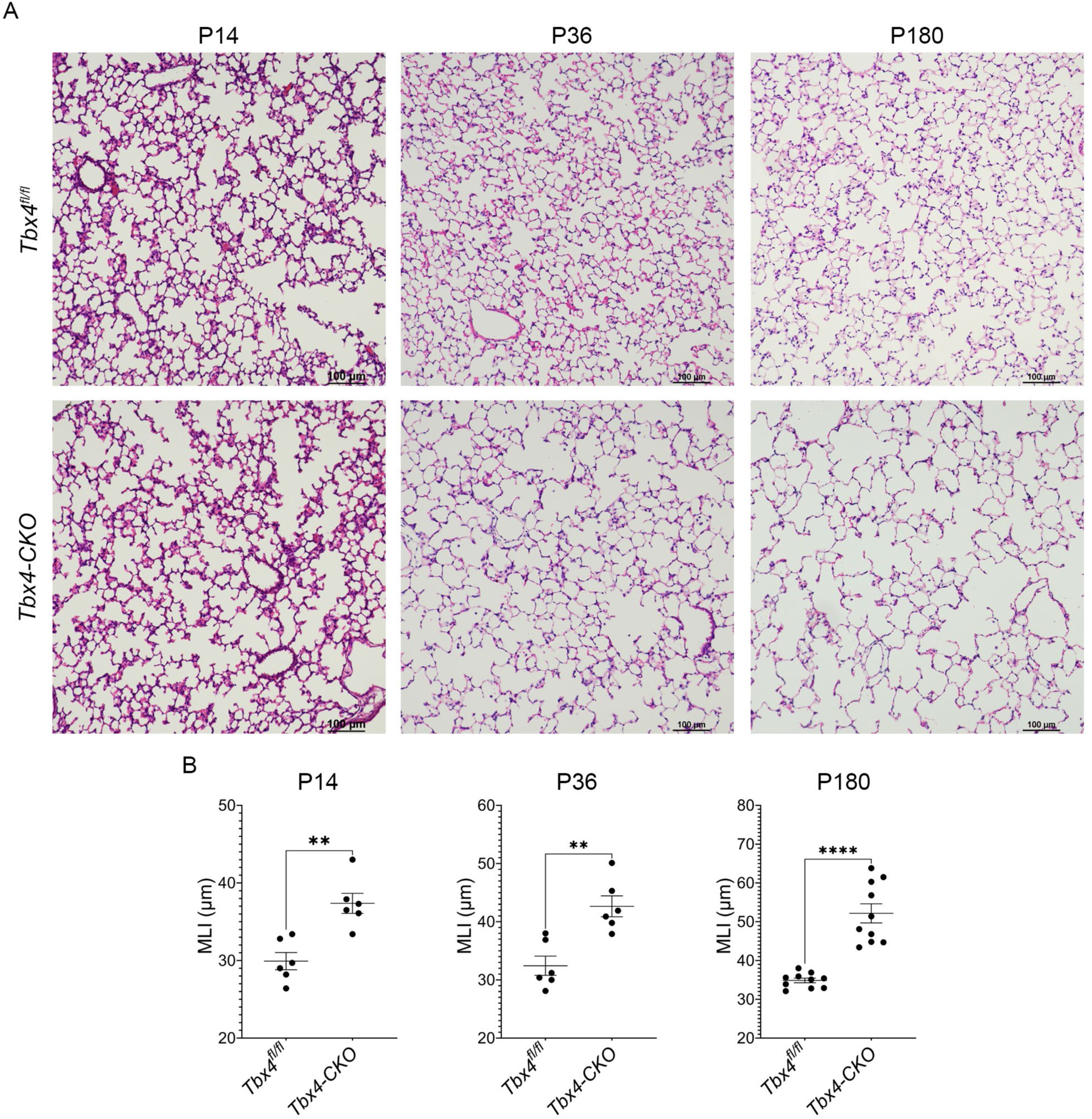
Disruption of *Tbx4* expression in the embryonic lung disrupts alveolar lung development. (A) Photomicrographs of H&E-stained lung sections from mice of the indicated postnatal day. The top and bottom rows show the photos from the control and *Tbx4-CKO* lungs, respectively. (B) MLI calculations from lungs of the indicated timepoints. The graphs show individual values and mean ± SEM. Each dot represents the average MLI for a single mouse. The number of mice per group is as follows: n = 6 (P14); n = 6 (P36); n = 10 (P180). The statistical test used was a parametric unpaired two-tailed *t test*. \*\**P* < 0.01; \*\*\*\**P* < 0.0001.

Among the most relevant cell types for the integrity of the alveolar structure include alveolar type I (AT1), alveolar type II (AT2), and pulmonary endothelial cells (EC) [24]. Therefore, to investigate the cellular mechanism underlying these histological abnormalities, we performed RNA ISH using cell type-specific RNAscope probes to label these cells on six-month-old lung sections. Based on nuclei staining, the total number of cells in *Tbx4-CKO* lungs show a statistically significant reduction of 18.5% (226.5 ± 5.87 versus 184.5 ± 15.99; *P* = 0.0487) (Figure E4). Similarly, the average number of *Pdpn*, *Sftpc*, and *Pecam-1*-positive cells is reduced in these lungs by 20.8% (15.63 ± 0.94 versus 12.38 ± 2.37; *P* = 0.2499), 18.8% (40.93 ± 0.47 versus 33.23 ± 1.07; *P* = 0.0021), and 22.8% (126.60 ± 8.78 versus 97.70 ± 11.23; *P* = 0.0894), respectively (Figure E4). However, the proportion of these cells is not different between the groups, which is probably due to the parallel reduction in the total number of cells and each of these cell types (Figure E4). Taken together, these results demonstrate that early disruption of *Tbx4* expression within the developing lung negatively impacts the latest stage of lung development, alveologenesis. However, the phenotype may not be severe to the extent of causing changes to the cellular composition.

### Postnatal *Tbx4-CKO* Lungs Show Pulmonary Vasculature Simplification

Alveologenesis and vascularization of the lung occur simultaneously during development, and disruption of one process may affect the other [23]. Since *Tbx4-CKO* mice develop alveolar simplification, we sought next to uncover whether pulmonary vascularization is affected in these lungs. To determine whether *Tbx4-CKO* lungs also show abnormal vascularization, we performed immunostaining using a vWF antibody to visualize positive vessels in six-month-old lungs. By simple examination of the tissue sections, we observed fewer vWF-positive vessels in *Tbx4-CKO* lungs (Figure 3A). To confirm this observation, we counted the number of vessels positive for vWF per HPF. Consistent with its adverse effects on lung alveolarization, early disruption of *Tbx4* expression in lung development results in a 39.3% (17.87 ± 0.81 versus 10.85 ± 0.60; *P* < 0.0001) decrease in the number of vWF-positive vessels in the *Tbx4-CKO* group, suggesting that these lungs develop pulmonary vascular simplification (Figure 3B).

**Figure 3.**
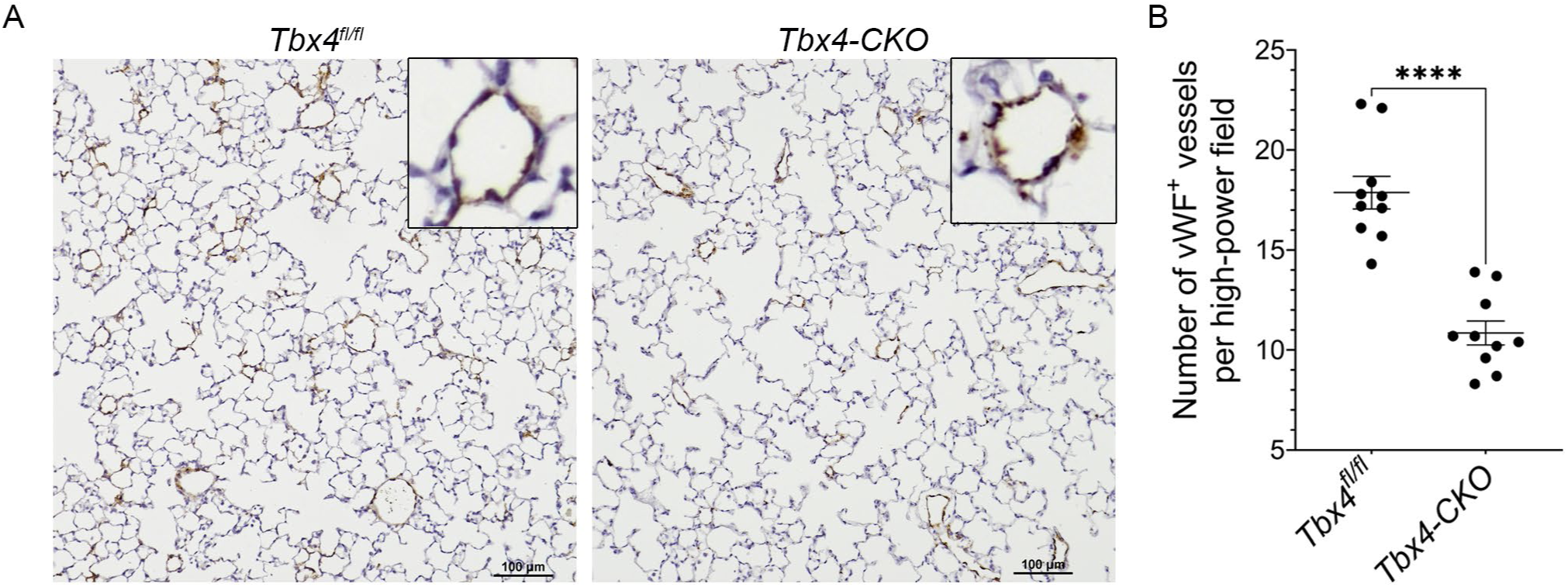
Six-month-old *Tbx4-CKO* lungs show pulmonary vascular simplification. (A) Representative photomicrographs of lung sections labeled with a vWF antibody using immunohistochemistry. Boxes in the top right corner of the photos, show a higher magnification of representative pulmonary arteries included in the analyses of this study. Scale bar, 100 μm. (B) Quantification of the number of vWF-positive vessels per high-powered-field. The graph shows individual values and mean ± SEM. Each dot represents the average number of stained vessels per mouse lung. The statistical test used was a parametric unpaired two-tailed *t test*. n = 10 per group; \*\*\*\**P* < 0.0001.

### Six-month-old *Tbx4-CKO* Mice Develop Physiological and Histological Features of Pulmonary Hypertension

The presence of alveolar simplification in *Tbx4-CKO* lungs suggests a reduction in the alveolar and capillary surface area, which may result in increased vascular resistance [25]. Therefore, we next sought to investigate whether early loss of *Tbx4* increases susceptibility to pulmonary hypertension. *Tbx4-CKO* mice and controls were housed under room air conditions, and RVSP was measured at six months of age. Compared to controls, *Tbx4-CKO* mice show a statistically significant increase of 15.7% (25.90 ± 0.41 versus 29.96 ± 0.60; *P* < 0.0001) in RVSP, consistent with the development of pulmonary hypertension (Figure 4A). Since sex-specific differences exist in the epidemiology of pulmonary hypertension, we examined the RVSP from males and females separately but did not observe differences beyond the females showing a more significant increase in RVSP (females: 25.25 ± 0.48 versus 30.23 ± 0.87; *P* < 0.0001 and males: 26.88 ± 0.61 versus 29.60 ± 0.81; *P* = 0.0125) (Figure 4A). Consistent with these results, *Tbx4-CKO* lungs show mild vascular remodeling, as confirmed by the calculation of the ratio of vessel wall thickness to the total area of the vessel, which shows a 14.2% (0.32 ± 0.005 versus 0.37 ± 0.011; *P* = 0.0013) increase (Figure 4B and 4C). Importantly, since the pressure overload in PAH can lead to right ventricular hypertrophy (RVH), we calculated the Fulton index (weight of the ventricle/(left ventricle+septum) and observed a median increase of 16.3% (0.21 ± 0.009 versus 0.25 ± 0.009; *P* = 0.0036), suggesting RVH (Figure 4D). Based on these results, we examined the function and structure of the right ventricle using echocardiography. *Tbx4-CKO* lungs show a lower TAPSE (0.86 ± 0.04 versus 0.76 ± 0.035; *P* = 0.0775), but it is not statistically significant (Figure E5). We also observed reduced right ventricular internal diameter (RVID) during systole (0.98 ± 0.05 versus 0.87 ± 0.03; *P* = 0.0568) and diastole (1.71 ± 0.07 versus 1.50 ± 0.03; *P* = 0.0097), but only the latter shows statistical significance (Figure E5). However, the fractional area change is not different between the groups (0.43 ± 0.02 versus 0.42 ± 0.02; *P* = 0.5605), suggesting that the RV function during systole is normal. Other relevant parameters, including cardiac output, stroke volume, and RV thickness, do not show significant differences between the groups (Figure E5). These results suggest that the PH phenotype is not severe enough to impair RV function in *Tbx4* mutant six-month-old mice.

**Figure 4.**
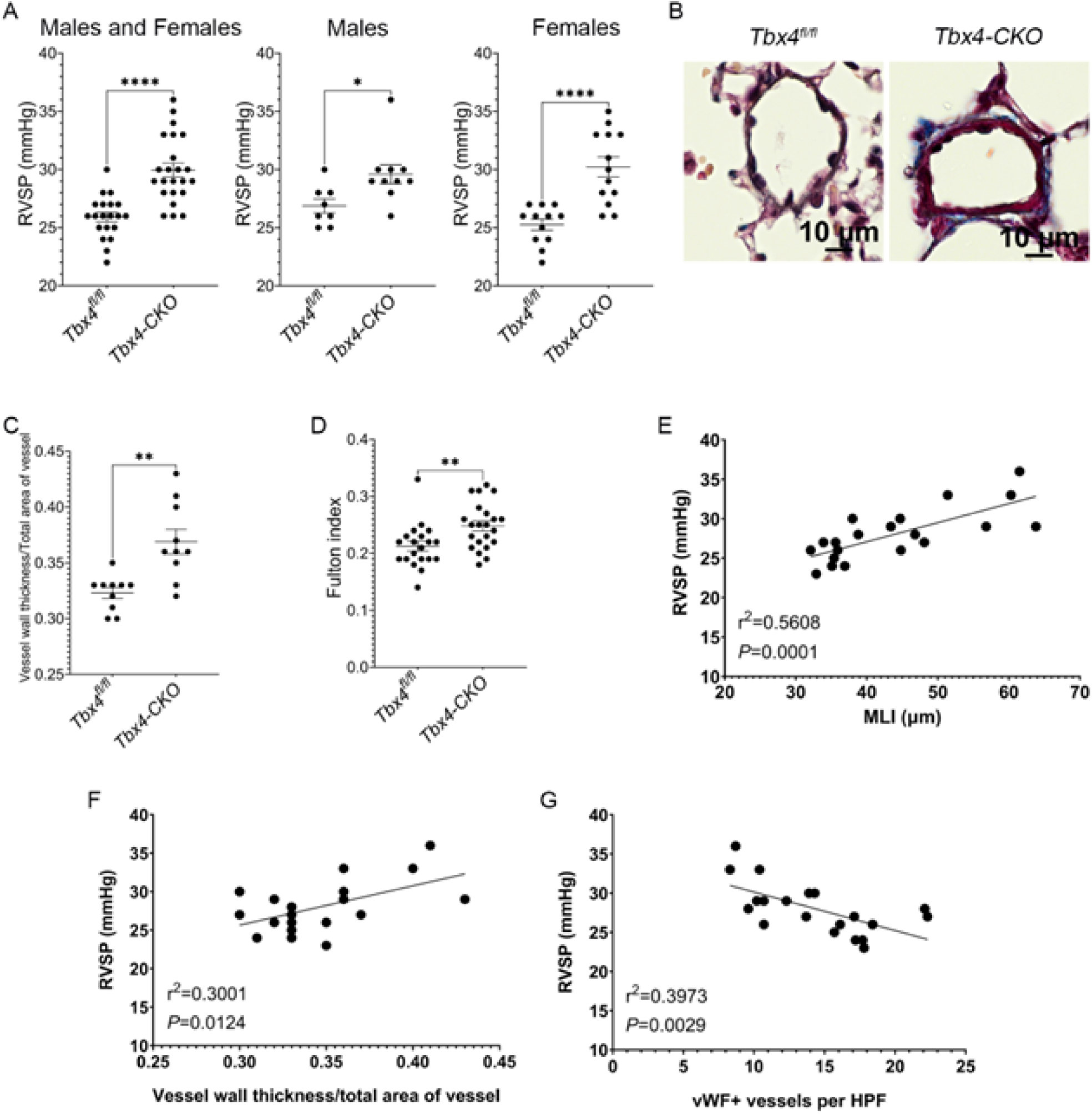
Physiological and histological features of pulmonary hypertension in six-month-old *Tbx4-CKO* mice. (A) RVSP measurements from six-month-old mice housed in room air. We evaluated differences in RVSP in a dataset with male and female values combined, but sex-specific differences were also assessed. The graphs show individual values and mean ± SEM. Each dot represents the RVSP for a single mouse. The number of mice per group is as follows: n = 20 (*Tbx4^fl/fl^*,M and F), n = 23 (*Tbx4-CKO*, M and F), n = 8 (*Tbx4^fl/fl^*,M), n = 10 (*Tbx4-CKO*, M), n = 12 (*Tbx4^fl/fl^*,F), n = 13 (*Tbx4-CKO*, F). (B) Representative photomicrographs of pulmonary arteries from six-month-old lung sections stained with Masson Trichrome and Verhoeff’s stain. (C) The ratio of vessel wall thickness to total area of the vessel was calculated to determine the degree of vascular remodeling. The graph shows individual values and mean ± SEM. Each dot represents the calculated ratio for a single mouse. n = 10 mice per group. (D) The Fulton index was calculated to evaluate for RVH. The graph shows individual values and mean ± SEM. Each dot represents the Fulton index for a single mouse. n = 20 (*Tbx4^fl/fl^*) and n = 22 (*Tbx4-CKO*). (E and F) Simple linear regression analysis showing how the MLI (E) and vessel wall thickness/total area ratio (F) influence the behavior of RVSP. The statistical test used was a parametric unpaired two-tailed *t test*. R-squared, r^2^; n = 10 (*Tbx4^fl/fl^*); n = 10 (*Tbx4-CKO*); \**P* < 0.05; \*\**P* < 0.01; \*\*\*\**P* < 0.0001.

To determine whether there is an association between the RVSP and the histological parameters measured in this study, we performed linear regression analyses. The results show a statistically significant positive linear relationship between RVSP and MLI (r^2^ = 0.5608, *P* = 0.0001) and RVSP and vessel wall thickness (r^2^ = 0.3001, *P* = 0.0124) (Figure 4E and 4F). Due to the enlarged airspaces in *Tbx4-CKO* lungs, a reduction in septal tissue density is expected, which may be accompanied by a decrease in pulmonary vascularization. Therefore, we also tested the relationship between MLI and the number of vWF+ vessels per HPF. We found a negative relationship between these two variables in *Tbx4-CKO* lungs (r^2^ = 0.6536, *P* < 0.00001) (Figure E6). To determine if the reduction in pulmonary vascularization influences RVSP, we compared RVSP and the number of vWF+ vessels per HPF and observed a negative relationship (r^2^ = 0.3973, *P* = 0.0029) (Figure E6). To a certain extent, these results demonstrate that enlargement of the airspaces, vascular simplification, and vascular remodeling contribute to the development of PH in *Tbx4-CKO* lungs, as expected.

### *Tbx4-CKO* Lungs Show Disruption of Essential Gene Regulatory Networks Relevant to Lung Alveolarization and Vascular Development

To understand the molecular mechanisms underlying abnormal alveologenesis and vascularization, we performed RNA sequencing analyses using P14 and P36 *Tbx4-CKO* lungs. Sequencing results show that there are 905 differentially expressed genes of which 663 are upregulated and 242 are downregulated in P14 lungs (Figure 5A). In P36 lungs, 418 out of 736 differentially expressed genes are upregulated, and the remaining 318 genes are downregulated (Figure 5B). A complete list of genes is provided in the online data supplement (File E1). Pathway analysis of the downregulated genes in P14 lungs revealed pathways associated with lung development, function, and disease. Among the top ten affected pathways include endothelin-3/EDNRB signaling, nitric oxide signaling, and EnaC regulation (Figure 5C). These pathways are important in the context of vascular homeostasis, respiratory function, and alveolar epithelial function [26–28]. Interestingly, the canonical WNT signaling is the most affected pathway (Figure 5C). This is relevant to the abnormal alveolar development phenotype in *Tbx4* mutant lungs since this pathway promotes the expansion of a subpopulation of *Axin2*-positive AT2 cells, an essential process during alveologenesis [29]. Closer examination of the differentially expressed genes in P14 lungs shows differential expression of genes encoding components of the WNT signaling pathway (Figure 5A). Of the downregulated genes, *Lgr5*, *Wnt3a*, and *Tnc*, have been shown to play a role in alveolar development [30–32]. Particularly, deficiency of *Lgr5* and *Tnc* negatively impacts lung alveolarization [31, 32].

**Figure 5.**
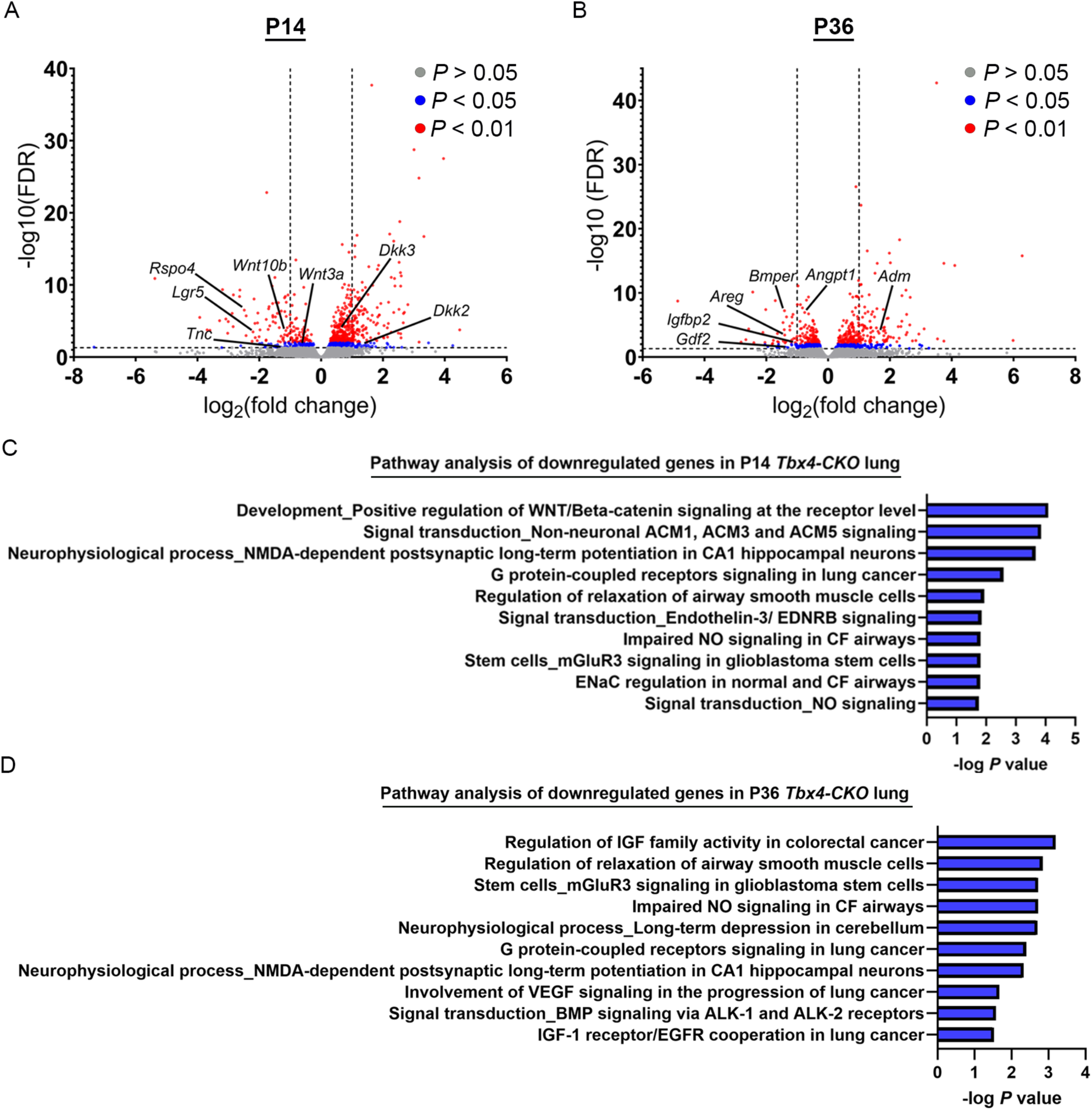
RNAseq analysis using P14 and P36 *Tbx4-CKO* lungs. (A and B) Volcano plot showing differential expression of genes in P14 (A) and P36 (B) lungs. The x-axis shows the direction of the fold change for each gene, and the y-axis is the -log10 of the adjusted *P* value (FDR). Each dot represents a single gene, and the color of the dots indicates their degree of significance, as described in the legend of the graph. The horizontal dashed lines indicate the cut-off for statistical significance (equivalent to 0.05), whereas the vertical lines define the threshold for fold change (equivalent to 2). Dots or genes on the left side of 0 are downregulated, and the ones to the right are upregulated. (C and D) Bar graphs showing the top ten affected regulatory pathways in P14 (C) and P36 (D) lungs, as determined by pathways analysis using MetaCore data. The x-axis shows the -log *P* value as calculated by MetaCore. The higher this number, the more statistically significant the pathway.

Relevant pathways to lung function and disease are affected in the P36 lungs as well, including the nitric oxide signaling (Figure 5D). The most affected pathway involves regulation of the insulin-like growth factors (IGF) family. Members of this family of growth factors have been described in the context of lung and vascular development, respiratory disease, fibrosis, and cancer [33–37]. The VEGF and BMP signaling pathways are also among the most affected in *Tbx4-CKO* lungs (Figure 5D). Consistent with these results, examination of the enriched genes in this dataset revealed differential expression of genes involved in lung development (*Adm* and *Igfbp2*), angiogenesis (*Adm, Angpt1, Areg, Bmper*, and *Igfbp2*), vascular disease (*Adm* and *Gdf2*), and fibrosis (*Igfbp2*) (Figure 5B) [38–44]. Surprisingly, we did not see differential expression of genes encoding components of the VEGF signaling pathway such as *Flt1*, *Flt4*, *Kdr*, *Vegfa*, *Vegfb*, and *Vegfc*. Taken together, the combined dysregulation of these genes may contribute to the various histological as well as PH phenotype observed in *Tbx4-CKO* lungs.

## Discussion

Human cases with *TBX4* mutations display a wide range of lung pathologic features, ranging from lethal lung developmental disorders to adult-onset pulmonary hypertension. Some pathologic features associated with *TBX4* variants include lack of acinar development, thickened interstitium, alveolar simplification, abnormal vascular development, vascular remodeling, and fibrosis [11]. In this study, we found that the mutant mouse lungs show an increasing enlargement of the airspaces over three different timepoints, demonstrating that TBX4 is essential for alveolar development. In addition, mutant adult mouse lungs contain regions with variable degrees of tissue thickening and collagen fiber deposition. In *Tbx4-CKO* mice, the extent of vascular remodeling is mild relative to the severity of the observations made in human cases. This may be due to mouse models not being prone to developing pulmonary vascular remodeling in general. These findings suggest that the combination of the histological abnormalities observed contributes to increased vascular resistance and the development of PH in these mice. Our results demonstrate that a reduction in *Tbx4* expression within the embryonic lung is sufficient to disrupt development and function and recapitulate phenotypes of human cases with *TBX4* mutations and PH. Importantly, our results support the idea that disruption of *Tbx4* expression is consistent with the etiology of not only group 1 PH but also group 3. Furthermore, the results demonstrate overlapping genetic mechanisms in the etiology of these subgroups of PH, similar to humans [45].

This is the first mouse study showing that disruption of *Tbx4* expression negatively impacts postnatal lung development and increases susceptibility to pulmonary hypertension. *Tbx4* is robustly expressed within the lung before birth with its peak mRNA expression at E15 [8]. A previous study shows that the *Tbx4* lung enhancer in our mouse model is active in lung mesenchymal cells, and that these cells give rise to fibroblasts, pericytes, smooth muscle cells, and endothelial cells [7]. Importantly, the study shows that the enhancer remains active at E18.5 in airway smooth muscle cells, and possibly, mesenchymal cells. Therefore, future studies using this mouse model are required to better characterize the effect of the loss of *Tbx4* at the canalicular, saccular, and early alveolar development.

Under normoxic conditions, *Tbx4-CKO* mice develop multiple histological abnormalities that may influence vascular resistance and lead to PH. For instance, the combination of alveolar simplification and reduced vascularization can reduce the pulmonary capillary bed, leading to an increase in pulmonary vascular resistance and hypertensive vascular changes [25]. These vascular changes can include intimal proliferation, vascular remodeling, medial hypertrophy, plexiform lesions, thrombosis, and vasoconstriction [46]. The consequences of these abnormalities include vascular stiffening, increased pulmonary vascular resistance, disturbed blood flow, and increased pulmonary arterial pressure [47]. The latter can also be caused by alveolar septal thickening due to fibrosis, which can stiffen the lung and prevent the vasculature from expanding to efficiently circulate blood. Notably, future studies should aim to identify which of these factors is the main driver of increased arterial pressure in *Tbx4-CKO* lungs. One step towards that goal may include determining whether elevated RVSP can be detected in early stages of postnatal lung development. The presence of elevated pressures in younger mice, which lack signs of vascular remodeling (Figure E7), would strongly suggest that the lung developmental abnormalities in *Tbx4-CKO* mice are a major factor contributing to PH.

Studying the loss of *Tbx4* expression in the context of hypoxia will also be of interest. Exposure to hypoxia may exacerbate the phenotypes observed in *Tbx4* mutant mice. The expectation is to observe fibrosis and signs of vascular remodeling before six months of age. In addition, in contrast to the six-month timepoint, we expect to see changes in the behavior of AT1 and AT2 cells in younger lungs, including apoptosis and proliferation. Needless to say, we also anticipate the detection of elevated RVSP at a younger age.

The molecular mechanisms described in this study provide a potential mechanism underlying the phenotypes in *Tbx4-CKO* lungs. The observed combination of gene expression changes in the P14 *Tbx4* mutant lungs suggests that the WNT signaling pathway is disrupted. At this stage, this pathway stimulates the expansion of a subpopulation of AT2 cells marked by *Axin2*, which are important for alveolar growth and maturation based on ex vivo organoid analyses [29]. Interestingly, the P36 RNAseq dataset shows downregulation of *Igfbp2*. Using mouse models of lung injury, it has been shown that reduced protein expression of *Igfbp2* increases markers of senescence in AT2 cells [48]. AT2 cells respond to lung injury by proliferating and replacing dead AT1 cells [49]. Therefore, these results may suggest that AT2 cells are less proliferative in *Tbx4-CKO* lungs, which can impair its repair process. *Igfbp2* also marks a subpopulation of AT1 cells that gradually increases during alveologenesis and accounts for approximately 95% of these cells by P60 [50]. If IGFBP2 plays important roles in the homeostasis of AT1 cells, its reduced expression may negatively impact their behavior in *Tbx4* mutant lungs. Based on this evidence, we are surprised to see that the expression of gene markers of AT1 and AT2 cells is not significantly reduced in our RNAseq datasets. This suggests that the disruption of these pathways is not sufficient to affect alveolar epithelial cell behavior and that parallel pathways may also play a role in their development and homeostasis. Nevertheless, the disrupted pathways may be essential for the growth and function of other lung cell types, including fibroblasts and pericytes, which can also contribute to the phenotypes observed in this study.

Of the differentially expressed genes in the P14 RNAseq dataset, *Tnc* and *Lgr5* deserve particular attention. The former is an extracellular matrix glycoprotein composed of six monomers with an average size of 215 kilodaltons [51]. TNC plays an essential role in lung development and is detected robustly in the lung mesenchyme and tip of forming secondary septa during alveolarization [52]. Knocking out *Tnc* delays alveolarization and impairs microvascular maturation in the early stages, but the phenotype is resolved [31]. Regarding *Lgr5*, it encodes a G-protein coupled receptor and has been reported to be a marker of a subpopulation of lung mesenchymal cells [30]. Mice lacking a copy of *Lgr5 show* histological features suggestive of abnormal alveolar development [32]. Although the phenotype observed in *Tbx4-CKO* mice does not exactly recapitulate the findings from *Tnc*-knockout or *Lgr5−/+* mice, the downregulation of these genes, as well as others involved in alveologenesis, may have a combinatorial effect that contributes to abnormal alveolar development in the mice reported in this study.

As described above, six-month-old *Tbx4-CKO* mice show reduced vascularization. Interestingly, endothelial cell composition is normal in the mutant lungs at that age, suggesting that loss of vasculature due to cell apoptosis does not underlie that phenotype. In addition, these findings suggest that the vascularization phenotype may have a developmental origin. The RNAseq dataset from P36 lungs show enrichment for genes relevant to angiogenesis, including *Bmper*, *Areg*, *Adm*, *Igfbp2*, and *Angpt1* (fold change = −1.7) (Figure 5E) [39, 40, 42–44]. *In vitro* downregulation of *Bmper* and *Areg* leads to reduced endothelial cell tube formation [39, 40]. Since these are pro-angiogenic factors, their downregulation in *Tbx4-CKO* lungs may have a combinatorial effect impairing angiogenesis, and as a consequence, causing abnormal vascular development. Further gene silencing experiments are required to investigate the contribution of *Bmper* and *Areg* to abnormal development in *Tbx4* mutant lungs.

It is important to emphasize that we did not examine gene expression changes at the single-cell resolution. Single-cell RNA sequencing analyses are required to study the differential expression of genes in the context of specific cell types. This approach would provide insights into how changes in gene expression profiles and representation of specific lung cell populations contribute to the phenotypes observed in this study. In addition, it would help identify potential targets to perform rescue experiments. Lastly, future studies should focus on developing approaches to restore the expression of *Tbx4* in *Tbx4-CKO* lungs.

## Acknowledgements

We thank the following fee-for-service cores for their help with tissue sectioning, imaging and RNAseq analysis, respectively: Histology Core of the Indiana Center for Musculoskeletal Health, the Indiana Center for Biological Microscopy, and the Center for Medical Genomics (CMG). These cores are supported by the IU School of Medicine and the Indiana Clinical Translational Sciences Institute (CTSI). The CMG is partially supported by the Indiana University Grand Challenges Precision Health Initiative.

