## Supplementary material for "Loss of *Tbx4* Affects Postnatal Lung Development and Predisposes to Pulmonary Hypertension": Online data supplement


| **Table of Contents** | **Pages** |
| --- | --- |
| Supplemental Methods | 2-9 |
| Supplemental References | 10 |
| Supplemental Tables | 11 |
| Supplemental Figures | 12-18 |

### Supplemental Methods

#### Mouse Models and Genotyping

The *Tbx4*$^{LME}$*-Cre* (stock no. 033331) and *Tbx4*$^{cond}$ (stock no. 043812-JAX) mouse models were purchased from The Jackson Laboratory [1, 2]. *Tbx4*$^{LME}$*-Cre* and homozygous *Tbx4*$^{cond}$ mice were crossed to generate the *Tbx4* conditional knockout (*Tbx4-CKO*) mice used in this study. *Tbx4*$^{fl/fl}$ mice served as littermate control animals. Genotyping was carried out by regular polymerase chain reaction (PCR) using the primer sequences recommended by the vendor (Table E1). All animal maintenance and procedures were performed in accordance with the Animal Care and Use Committee at Indiana University School of Medicine, protocol 20135.

#### Harvesting of Mouse Embryos

Pregnancy was determined by the presence of a vaginal plug. Pregnant dams were euthanized at the appropriate gestational day to harvest embryos of the desired embryonic age. Embryos were removed from the uterus, followed by removal of the placenta, yolk sac, and amnion. The yolk sac or tail of the embryo was used for DNA extraction and subsequent genotyping analyses. The embryos or their lungs were fixed in 4% paraformaldehyde overnight at 4 ̊C. After fixation, the tissues were washed with 1X PBS and underwent a series of ethanol dehydration steps. Whole embryos or their lungs used for RNA in-situ hybridization (ISH) were stored in absolute ethanol at -20 ̊C until the experiment was carried out. The embryos meant for sectioning, after overnight fixation at 4 ̊C, went through a series of ethanol dehydration steps followed by paraffin embedding in transverse position.

#### Harvesting and Processing of Mouse Lung Tissues

Mice at various postnatal stages were euthanized and lungs were inflated by instillation with 10% neutral buffered formalin (NBF) using a tracheal cannula at a constant pressure of 20 or 25 cm$H_2O$ [3]. The cannula was 22 gauge for P14 and 20 gauge for P36 and six-month-old lungs. Once the post-caval lobe expanded and no visible expansion of the lungs was observed, the lungs were considered fully inflated. The trachea was ligated with a suture to preserve the pressure. The lungs, along with the heart and thymus, were removed *en bloc* and submerged in 10% NBF for approximately 24 hours at 4 ̊C. After fixation, the lungs underwent dehydration steps with ethanol, tissue clearing with xylene, and paraffin embedding. The P14 lungs were embedded as a whole and sectioned at a 7 μm thickness from the dorsal surface of the lungs to get uniform sectioning of all the lobes. The P36 and six-month-old lungs were separated by ligating the primary bronchus of the left lung with a suture to keep the pressure inside the lung. The left lung was then cut in two locations to get three pieces, which were processed and embedded individually [4]. The individual pieces were cut into 5 μm thick sections following the longitudinal axis of the lung.

**Histological Analysis of the Lung**

While carrying out the analyses of lung histology, the researcher was unaware of the experimental condition of each lung when possible. All the photos were taken with a Nikon Ti2E Fully Automated Live Cell Inverted Microscope unless otherwise indicated. For mean linear intercept (MLI) and pulmonary vascularization analyses, we focused on the distal areas of the lung while avoiding large airways and vessels when possible. MLI assessment was performed using serial lung tissue sections stained with hematoxylin and eosin (H&E) stain. Photographs from up to 5 random non-overlapping regions of the left lung were taken at 20X magnification. Subsequently, the photographs were viewed under a field of equidistant horizontal lines dividing it into four segments of the same size. The MLI was calculated by dividing the total length of a line by the number of intercepts of tissue [5]. After calculating the MLI from all photographs of a single lung, the average was calculated and used as the final value in this study. Immunohistochemical staining for the endothelial marker von Willebrand factor was used to assess pulmonary vascularization of six-month-old lungs. The number of vWF-positive vessels with a diameter smaller than 100 μm in a field with original magnification of 200X were counted and expressed as high-power field [6]. To examine the vessel wall thickness, the tissue sections were rehydrated and stained with Masson's Trichrome and Verhoeff stain to easily visualize the elastin in the blood vessels. Subsequently, the slides were imaged at 40X magnification using the Aperio Scanscope Imaging System. To determine the pulmonary vessel wall thickness of intra-acinar blood vessels, the inside vessel wall area was subtracted from the outer vessel wall area and divided by the outer wall area to obtain the ratio of vessel thickness to total area of vessel. Only vessels that were 25-100 μm in diameter were included in the analysis. Twenty vessels per animal were measured.

**Echocardiography**

Mice were anesthetized with 5% Isoflurane in an induction box, placed on a heated platform with a nose cone, and maintained at 1-2%. The hair was clipped over the chest, and fine hair was removed using a depilatory cream. The skin was cleaned with wet gauze squares, and subsequently, ultrasonic gel was placed over the chest for the echo procedure, and an ultrasonic probe was placed in contact with the gel. The probe was moved to acquire different images in M-Mode, B-Mode, and PW doppler. When finished, the chest was cleansed with gauze squares, and the mouse was returned to its cage. Measurements were taken using a Vevo 2100 ultrasound machine (VisualSonic) with a 40 MHz probe. The Vevo Lab software was used to measure diastolic and systolic dimensions, RVOT, TAPSE, cardiac output, stroke volume, pulmonary acceleration time (PAT), and pulmonary ejection time (PET).

**Right Heart Catheterization**

Mice were anesthetized by inhalation of 4-5% isoflurane in an induction box and placed on a nose cone with Isoflurane maintained at 2%. The mouse was placed on a heating pad to keep the body temperature of the animal at approximately 37 ̊C. A midline neck incision of about 3 cm was made and the right external jugular vein was cannulated with a 1.4F Millar catheter, which was advanced into the right ventricle. Once the rodent was stabilized, RVSP data was collected, and the catheter was removed. The carotid artery was isolated, and the same 1.4F catheter was inserted in the aorta to measure systemic pressure. Subsequently, the catheter is advanced into the left ventricle. Once the rodent stabilized, LVSP data was collected, and the catheter was removed. At the end of the procedure, the anesthetized mouse was sacrificed via exsanguination and excision of the heart. The measurements took approximately 10 minutes per animal.

**RNA *In situ* Hybridization (ISH)**

A previously published whole-mount or tissue sections RNA ISH protocol was used to evaluate gene expression in mouse embryos [7, 8]. Briefly, plasmids with the *Tbx4* and Cre recombinase cDNA sequences were purchased from Genscript (clone ID: Omu08335D) and Sigma Aldrich (Cat. No. OGS591), respectively. The *Tbx4* ORF was cloned into a pcDNA3.1+/C-(K)-DYK vector and the Cre recombinase into a pSF-CMV vector. The DNA fragment was amplified from cDNA using PCR and the primers indicated in Table E1. The fragment was purified using DNA gel purification reagents (Qiagen) and subsequently used as the template for an *in vitro* transcription reaction using the DIG RNA Labeling Kit (SP6/T7) (Sigma Aldrich, Cat. No. 11175025910). After riboprobe synthesis, the whole embryos or embryonic lungs stored at -20 ̊C in 100% ethanol were allowed to equilibrate to room temperature for one hour. Subsequently, they were rehydrated with graded methanol series, permeabilized with proteinase K (10 μg/mL) at room temperature for 10 minutes (E10.5 embryos) or 5 minutes (E11.5 lungs), and pre-hybridized for 1 h at 70 ̊C with slow rotation. Hybridization took place overnight at 70 ̊C. After post-hybridization washes, the tissues were blocked at room temperature for 2 h with 2% blocking reagent (Sigma Aldrich, Cat. No. 11096176001). Blocking solution was replaced with anti-digoxigenin antibody diluted (1:5000) in 2% blocking reagent and incubated overnight at 4 ̊C. Color development was performed with BM purple AP substrate (Sigma Aldrich, Cat. No. 11442074001) at room temperature and took approximately 3 h. For the paraffin sections, multiple optimizations were made to the protocol. The slides were heated at 60 ̊C for 30 minutes before deparaffinization with xylene and ethanol washes. Subsequently, the slides were taken through multiple washes as follows: three washes with xylene (10 minutes each), two 100% ethanol (5 minutes each), 90% ethanol (5 minutes), 70% ethanol (2 minutes), 50% ethanol (2 minutes), 30% ethanol (2 minutes), two 1X PBS (2 minutes each). Slides were then fixed with 4% paraformaldehyde (PFA) for 20 minutes at room temperature and washed twice with 1X PBS for 5 minutes each time. After fixation, slides were permeabilized with protein K diluted (1:5000 from a 20 mg/mL stock solution) in 1X PBS for 4 minutes at room temperature, washed in 0.2% glycine/1X PBS solution for 1 minute, and washed twice in 1X PBS for 1 minute each. Slide were fixed again with 4% PFA for 10 minutes at room temperature, followed by two washes in 1X PBS (5 minutes each) and water rinse. Before hybridization, the slides were treated with a solution (pH = 7) composed of 0.1 M triethanolamine (TEA) (Sigma Aldrich, Cat. No. 90279-100ML) and acetic anhydride (1:400) (Sigma Aldrich, Cat. No. 539996-25G) twice for 10 minutes each time. Subsequently, slides were washed twice in 1X PBS (5 minutes each), dehydrated (30%, 50%, 70%, 90%, and four 100% ethanol washes for two minutes each), and dried for 1 h at 65 ̊C. Slides were submerged in the hybridization solution with the corresponding riboprobe and incubated overnight at 70 ̊C. After hybridization, the slides were washed as described previously [7]. During the antibody treatment, the slides were submerged in 2% blocking reagent for 2 h at room temperature. After the blocking step, the blocking solution was replaced with an anti-digoxigenin/2% blocking solution, and the slides were incubated overnight at room temperature. Color development was performed following the previously published protocol.

**RNAscope Assays and Cell Quantification**

The RNAscope Multiplex Fluorescent Reagents Kit v2 (Advanced Cell Diagnostics, Intro Pack, Cat. No. 323136) was used following the manufacturer's instructions, specifically, the procedures describing the fixed formalin paraffin embedded protocol (Doc. No. UM 323100, revision B). The following probes were used: *Pdpn* (AT1 cell marker, Cat. No. 437771); *Sftpc* (AT2 cell marker,

Cat. No. 314101); and *Pecam1* (endothelial cell marker, Cat. No. 316721). Recommended positive and negative control probes provided by the kit were used to evaluate RNA integrity and specificity. The various probes were visualized using the following opal fluorophores at 1:1500: Opal 520 (detected under FITC optics, Akoya Biosciences Cat. No. FP1487001KT); Opal 570 (detected under Cy3 optics, Akoya Biosciences Cat. No. FP1488001KT); and Opal 690 (detected under Cy5 optics, Akoya Biosciences Cat. No. FP1497001KT). Nuclei were counterstained using DAPI stain supplied by the kit. Subsequently, a coverslip (Fisher Scientific Cat. No. 12-541-033) was applied using the ProLong Gold Antifade Mountant reagent (Thermo Fisher Scientific, Cat. No. P36930) and dried overnight at room temperature.

Upon completion of the RNAscope assay, tissue sections were photographed with a 40X objective on a Leica TCS SP8 Resonant-scanning or Leica TCS SP8 Dive confocal microscope. *Pdpn*-, *Sftpc*-, and *Pecam1*-positive cells from paraffin-embedded RNAscope confocal photos taken with a 40X objective were counted using the QuPath software version 0.4.4 [9, 10]. The technical note provided by Advanced Cell Diagnostics (Doc. No. MK 51-154, revision A) was used to operate the QuPath software. Cell counting for each probe was as follows: an average of 54 *Pdpn* (n=4 $Tbx4^{fl/fl}$ and n=4 *Tbx4-CKO* mice; four photos per mouse), 146 *Sftpc* (n=3 $Tbx4^{fl/fl}$ and n=4 *Tbx4-CKO* mice; four photos per mouse), and 434 *Pecam1* (n=4 $Tbx4^{fl/fl}$ and n=4 *Tbx4-CKO* mice; four photos per mouse) cells were counted per mouse. The percentage of cells positive for a specific probe was calculated as the ratio between probe-positive cells and total counted nuclei stained with DAPI in each photo multiplied by a hundred.

**Immunohistochemistry**

Lung sections from six-month-old mice were rehydrated and stained for the endothelial marker von Willebrand Factor (Biocare Medical, Pacheco,CA ; rabbit polyclonal anti-human vWF; CP039) following the manufacturer's instructions. Briefly, the tissue sections underwent a peroxide block (Peroxidazed 1; PX968), pepsin digest (CarezymeII: Pepsin; PEP956), and a protein block

(Rodent Block M; RBM961). Subsequently, the sections were incubated with the primary antibody at a dilution of 1:500 in Da Vinci Green (PD900) overnight at 4°C. After the incubation, tissue sections were treated with the Rabbit-on-Rodent HRP Poylmer (RMR961) and diaminobenzidine chromogen (Betazoid BDB2004). Lastly, sections were counterstained with Gills Hematoxylin (Vector Laboratories, Inc, Burlingame,CA; H-3401) and used for lung vascularization analyses.

**Bulk RNA sequencing**

Total RNA isolation from *Tbx4-CKO* and *Tbx4$^{fl/fl}$* lungs was performed using the miRNeasy Mini Kit (Qiagen, Cat. No. 217004). The RNA integrity number (RIN) for each sample was obtained using a 4200 TapeStation system (Agilent). Samples with a RIN of 5 or higher were included in the experiment. Library preparation was performed using the KAPA RNA HyperPrep Kit (Roche) and total RNA. Paired-end sequencing with read length of 100 bp was performed on an Illumina NovaSeq 6000 instrument.

The quality of the raw sequence was assessed with FastQC (Babraham Bioinformatics, Cambridge, UK) and no trimming was required. Subsequently, the sequencing reads were mapped to the UCSC mm10 reference genome using Spliced Transcripts Alignment to a Reference (STAR) software version 2.7.10a, and the quality of read mapping was determined with bamUtils (NGSUtilsJ/bam-filter/bam-stats, version 0.4.17) [11, 12]. The percentage of uniquely mapped reads involved a minimum of 88.9%, median of 90.8%, and maximum of 92.2%. After exclusion of low quality mapped reads, featureCounts version 2.0.3 was used to quantify gene expression levels [13]. Differential expression analysis was performed using edgeR version 3.38.4 [14]. Genes with a false discovery rate (FDR) $< 0.05$ and fold-change $> 2$ were considered significant. A list of genes that met these criteria was uploaded to the MetaCore software (Source: MetaCore™, a Cortellis solution, 04/07/2023, © 2023 Clarivate™) to carry out pathway analyses. Sequencing analysis was carried out in the Center for Medical Genomics at Indiana University

School of Medicine, which is partially supported by the Indiana University Grand Challenges Precision Health Initiative.

**Quantitative Reverse Transcription Polymerase Chain Reaction (RT-qPCR)**

For targeted gene expression analyses, we performed RT-qPCR using total RNA. First, 500 ng of RNA was used in reverse transcription reactions using SuperScript IV Reverse Transcriptase (Life Technologies, Cat. No. 18090200). Real time PCR was performed using the QuantiTect SYBR Green PCR kit (Qiagen, Cat. No. 204145) and the LightCycler 480 II (Roche). Relative mRNA expression was quantified using the $\triangle\triangle C_t$ method. Expression levels of target mRNA were normalized to the expression of *Gapdh*. The primers used in this experiment are listed in Table E1.

**Statistical Analysis**

To assess the normality of the data, we used the Shapiro-Wilk test. Subsequently, a parametric unpaired two-tailed *t test* or nonparametric Mann-Whitney test was used to evaluate the statistical significance of the difference between the experimental groups. To evaluate for a direct relationship between variables, a simple linear regression analysis was performed using ten mice per experimental group. The statistical significance was defined as $P < 0.05$. All statistical analyses were performed using GraphPad Prism version 10.0.2 (232) for Windows, GraphPad Software, Boston, Massachusetts USA, www.graphpad.com.

### Supplemental Tables

#### Table E1. Genotyping, RT-qPCR, and riboprobe template primers information.

| Primer name | Primer sequence (5' - 3') | Fragment size (bp) | Application | Accession No. | Notes |
|---|---|---|---|---|---|
| Tbx4Cond-F | TGCTTTTAGCATCAGCACTGT | ~400 bp (*Tbx4*$^{cond}$) 321 bp (WT) | Genotyping | | |
| Tbx4Cond-R | TACCCACTTGTCCCTCCAAC | ~400 bp (*Tbx4*$^{cond}$) 321 bp (WT) | Genotyping | | |
| Tbx4LMECre-F | GCAGCCTTCCAGAAGCAGAGC | 160 bp | Genotyping | | |
| Tbx4LMECre-R | AGGCAAATTTTGGTGTACGG | 160 bp | Genotyping | | |
| P53-F | GGTTAAACCCAGCTTGACCA | 271 bp | Genotyping | NM_011640 | Endogenous control |
| P53-R | GGAGGCAGAGACAGTTGGAG | 271 bp | Genotyping | NM_011640 | Endogenous control |
| Tbx4exon5-F | CCATGCCAGGAAGACTTTAT | 88 bp | RT-qPCR | NM_011536.3 | |
| Tbx4exon5-R | TTCAGCTTCTGGAAAGAGAC | 88 bp | RT-qPCR | NM_011536.3 | |
| musTbx4-c4F | CATACTTCTCATCGACATCGT | 351 bp | RT-qPCR | NM_011536.3 | |
| musTbx4-c6R | TAGGAGGTCACAGAGATGAA | 351 bp | RT-qPCR | NM_011536.3 | |
| musTBX4-c6F | TGACAGTGACCTGCGTGTG | 117 bp | RT-qPCR | NM_011536 | |
| musTBX4-c7R | GGGCTGACATCTGGCTTAG | 117 bp | RT-qPCR | NM_011536 | |
| mmus_GAPDH-c3F | GGTGCTGAGTATGTCGTGGA | 97 bp | RT-qPCR | NM_001289726.1 | |
| mmus_GAPDH-c4R | CGGAGATGATGACCCTTTTG | 97 bp | RT-qPCR | NM_001289726.1 | |
| Crerecombinase-F | TAAGCAAAGCTTCGACCAAGTGACAGCAATGC | 625 bp | riboprobe template | | Contains a HindIII restriction site |
| Crerecombinase-R | TAAGCAGGATCCCACCAGCTTGCATGATCTCC | 625 bp | riboprobe template | | Contains a BamHI restriction site |
| musTbx4-c499F | GATGTTCCCCAGCTACAAGGT | 416 bp | riboprobe template | NM_011536.3 | Anneals to exon 4 |
| musTbx4-c894R | TGTGGTTCTGGTAGGAGGTCA | 416 bp | riboprobe template | NM_011536.3 | Anneals to exon 6 |

**Supplemental Figures**

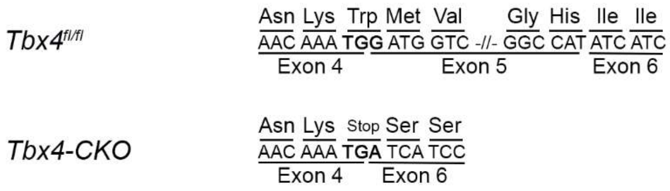


**Figure E1. Excision of *Tbx4* exon 5 creates a premature stop codon.** This figure illustrates a partial sequence of the wild-type *Tbx4* exon 4, exon 5, and exon 6. The sequence is divided into codons, and the encoded amino acid is shown on top. The *Tbx4$^{fl/fl}$* mice have a TGG codon at the junction of exons 4 and 5. Removal of exon 5 in *Tbx4-CKO* mice generates a TGA codon at the junction of exon 4 and 6, which is predicted to prematurely terminate translation and produce a truncated protein.

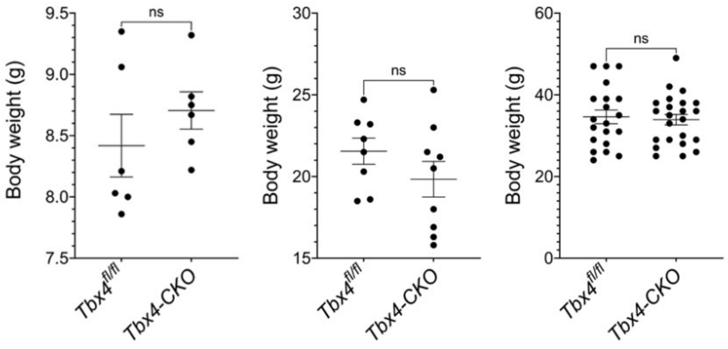


**Figure E2. *Tbx4-CKO* mice are healthy based on body weight measurements.** Body weight of P14 (left) (n = 6 per group), P36 (middle) (n = 8 *Tbx4*$^{fl/fl}$ and n = 9 *Tbx4-CKO*), and P180 (right) (n = 20 *Tbx4*$^{fl/fl}$ and n = 23 *Tbx4-CKO*) mice. The graphs show individual values and mean ± SEM. Each dot represents the body weight for a single mouse. n.s. = not significant.

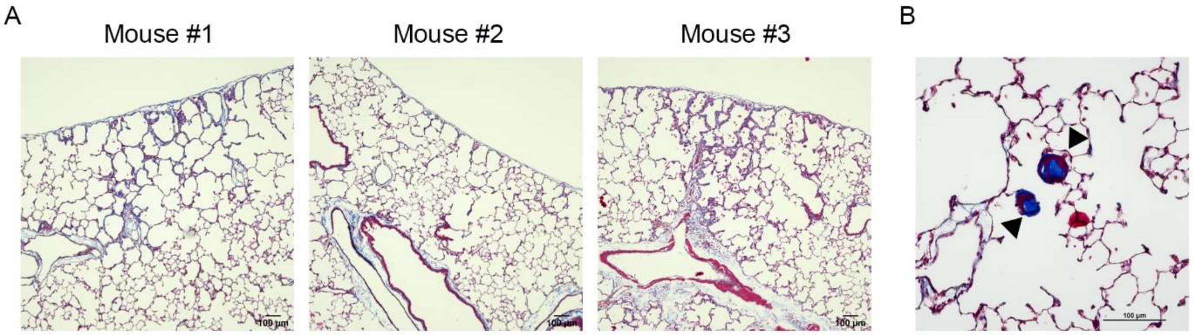


**Figure E3.** ***Tbx4-CKO*** **mice show tissue thickening, collagen fiber deposition, and organized thrombi.**

**Figure E3.** ***Tbx4-CKO*** **mice show tissue thickening, collagen fiber deposition, and organized thrombi.** (A) Three representative photomicrographs from six-month-old *Tbx4-CKO* lungs taken with a 10X objective. The lung sections were stained with Masson Trichrome and Verhoeff's stain. Tissue thickening and collagen fiber deposition (blue) are visible. (B) Photomicrograph from the six-month-old *Tbx4-CKO* lung that showed organized thrombi. The latter are indicated with black arrowheads.

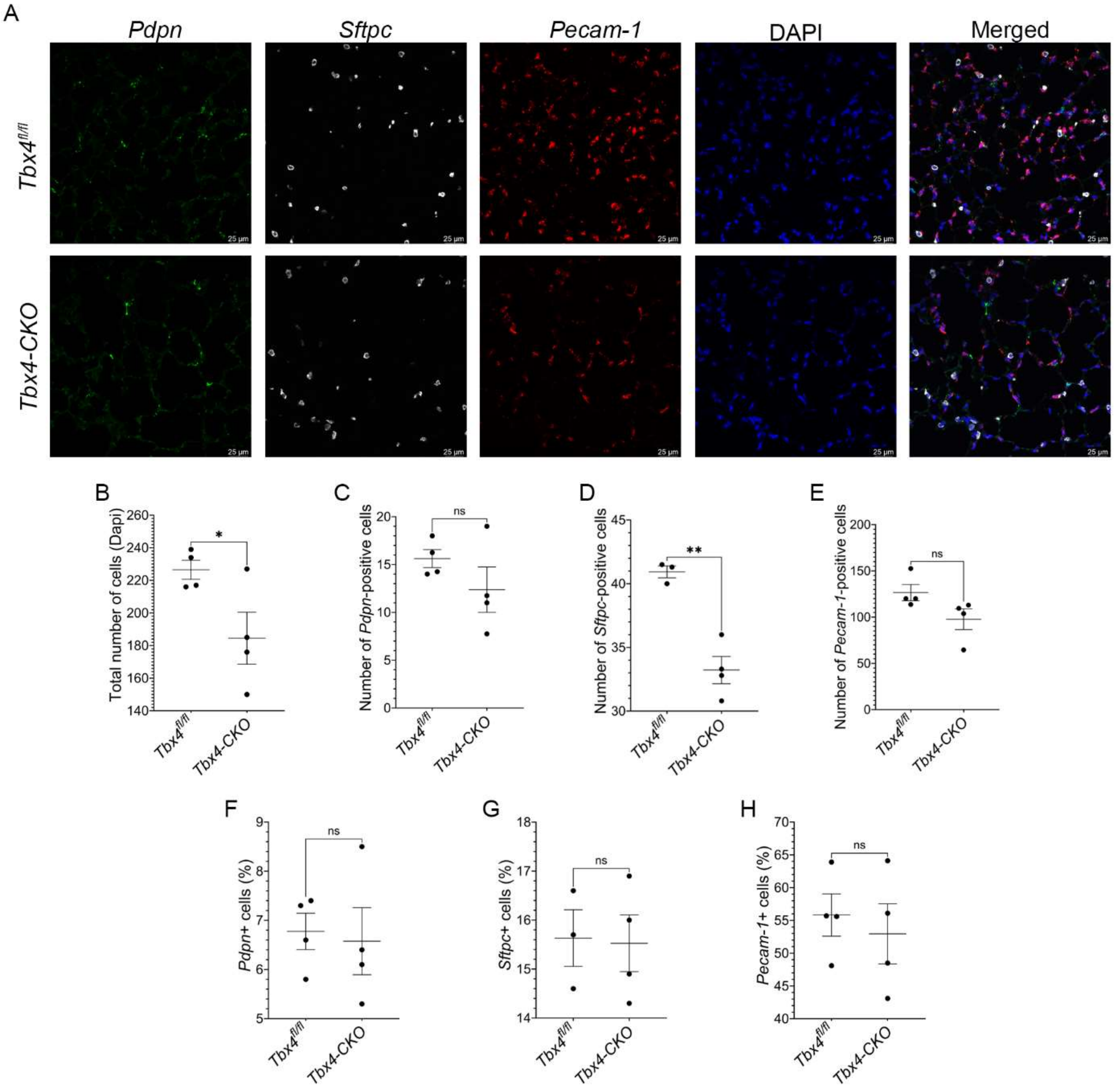


**Figure E4. Cell counting analysis of AT1, AT2, and endothelial cells in *Tbx4-CKO* lungs.** (A) Confocal images showing AT1 cells labeled with *Pdpn* (green), AT2 cells with *Sftpc* (white), and endothelial cells with *Pecam-1* (Red). Nuclei were identified with Dapi staining (blue). The last column shows a merged image with all the probes. (B) The total number of cells per 40X image were counted using nuclei-based counting (n = 4 per group). (C-E) The *Pdpn* (n = 4 per group) (C), *Sftpc* (n = 3 $Tbx4^{fl/fl}$ and n = 4 *Tbx4-CKO*) (D), and *Pecam-1*-positive cells (n = 4 per group) (E) were counted by identifying the probes overlapping with Dapi. (F-G) To evaluate changes in the percentage of cells, the ratio of *Pdpn* (n = 4 per group) (F), *Sftpc* (n = 3 $Tbx4^{fl/fl}$ and n = 4 *Tbx4-CKO*) (G), and *Pecam-1* (n = 4 per group) (H) to the total number of cells in the image was calculated and multiplied by 100. The graphs show individual values and mean ± SEM. Each dot represents the average number of cells or percentage for a single mouse. ***P* < 0.01; ns = not significant.

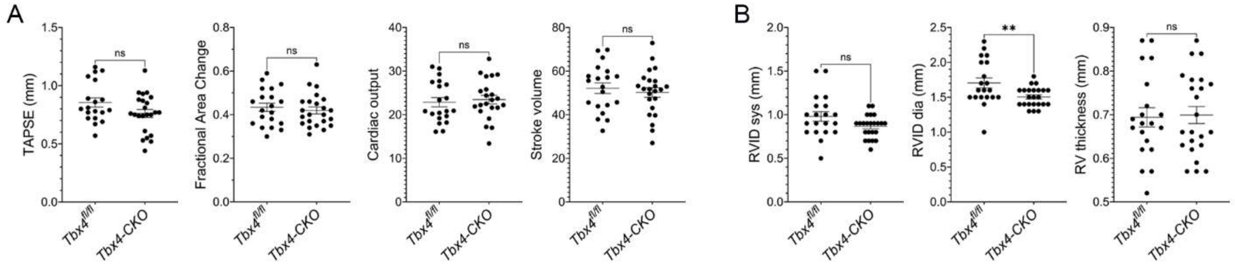


**Figure E5. Echocardiography parameters measured from six-month-old *Tbx4-CKO* mice.** (A) Echocardiography parameters used to assess RV function. (B) Echocardiography parameters to assess RV structure. The graphs show individual values and mean ± SEM. Each dot represents the parameter measured for a single mouse. n ≈ 20 mice per group. $^{**}P < 0.01$; ns = not significant.

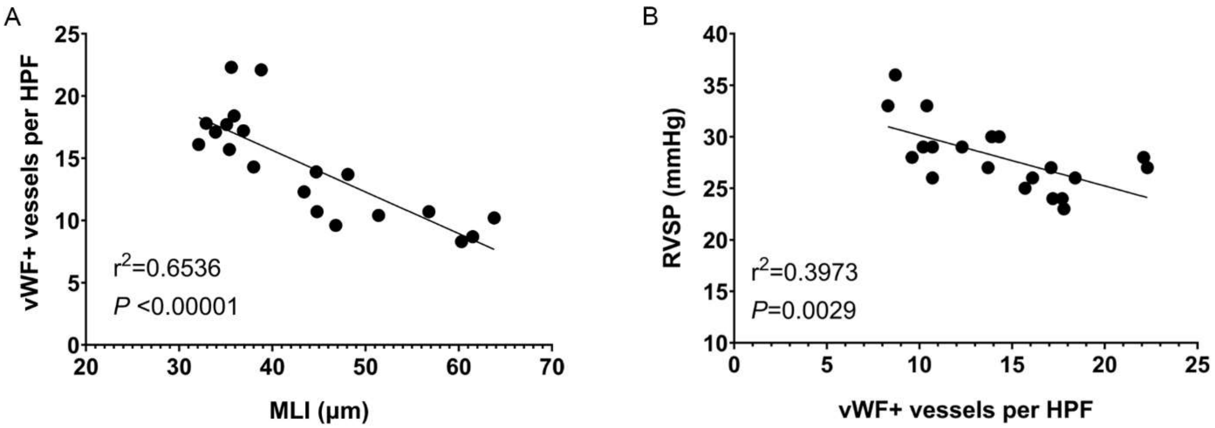


**Figure E6. Linear regression analysis comparing the number of vWF positive vessels with MLI and RVSP.** (A) Simple linear regression analysis showing how the MLI influences the number of vWF+ vessels per high power field. (B) Simple linear regression analysis showing how the number of vWF+ vessels per high power field influences the behavior of RVSP. R-squared, $r^2$; n = 10 (*Tbx4*$^{fl/fl}$); n = 10 (*Tbx4-CKO*); ***P* < 0.01; *****P* < 0.0001.

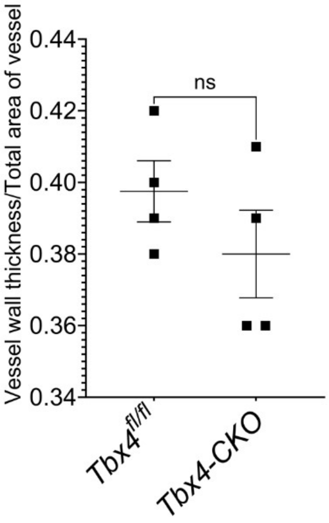


**Figure E7. P36 *Tbx4-CKO* lungs do not show vascular remodeling.** The ratio of vessel wall thickness to total area of the vessel was calculated to determine the degree of vascular remodeling. The graph shows individual values and mean ± SEM. Each dot represents the calculated ratio for a single mouse. n = 4 mice per group. ns = not significant.
